# Extracellular matrix viscoelasticity regulates mammary branching morphogenesis

**DOI:** 10.1101/2025.05.15.654249

**Authors:** Daniella I. Walter, Juliette W. Moore, Abhishek Sharma, Ryan S. Stowers

## Abstract

Structural and mechanical cues from the extracellular matrix (ECM) regulate tissue morphogenesis. Tissue development has conventionally been studied with *ex vivo* systems where mechanical properties of the extracellular environment are either poorly controlled in space and time, lack tunability, or do not mimic ECM mechanics. For these reasons, it remains unknown how matrix stress relaxation rate, a time-dependent mechanical property that influences several cellular processes, regulates mammary branching morphogenesis. Here, we systematically investigated the influence of matrix stress relaxation on mammary branching morphogenesis using 3D alginate-collagen matrices and spheroids of human mammary epithelial cells. Slow stress relaxing matrices promoted significantly greater branch formation compared to fast stress relaxing matrices. Branching in slow stress relaxing matrices was accompanied by local collagen fiber alignment, while collagen fibers remained randomly oriented in fast stress relaxing matrices. In slow stress relaxing matrices, branch formation was driven by intermittent pulling contractions applied to the local ECM at the tips of elongating branches, which was accompanied by an abundance of phosphorylated focal adhesion kinase (phospho-FAK) and β1 integrin at the tips of branches. On the contrary, we observed that growing spheroids in fast stress relaxing matrices applied isotropic pushing forces to the ECM. Pharmacological inhibition of both Rac1 and non-muscle myosin II prevented epithelial branch formation, regardless of matrix stress relaxation rate. Interestingly, restricting cellular expansion via increased osmotic pressure was sufficient to impede epithelial branching in slow stress relaxing matrices. This work highlights the importance of stress relaxation in regulating and directing mammary branch elongation.

## Introduction

Branching morphogenesis is a complex, multicellular process where cells self-organize into numerous types of glandular tissue, including mammary glands, lungs, and kidneys. This process is accompanied by remodeling of the extracellular matrix (ECM) architecture^1,2^. Much of our current understanding of branching morphogenesis comes from investigations into the effects of biochemical morphogens^2–5^ and transcriptional programs^6–8^, though recent work has revealed that the mechanics of the extracellular matrix can also regulate morphogenetic events^9–11^. Changes in ECM mechanics can influence a broad array of cellular processes, including differentiation, migration, proliferation, and morphological changes^12–14^. Within the context of tissue morphogenesis, properties of the ECM, such as matrix stiffness, fiber alignment, pore size, anisotropy, and ECM-bound cytokines, contribute to tissue patterning and organization throughout the developmental stages^15,16^. Despite this, delineating the structural, mechanical and molecular contributions of the ECM in guiding branch initiation and elongation within mammary epithelium remains challenging due to the dynamic and heterogeneous nature of the ECM.

Engineered extracellular matrices have been pivotal in isolating the role of mechanics and identifying processes that may guide mammary branching, as they enable precise and independent control of specific signals in both time and space, which is something that is difficult to achieve *in vivo*. For example, 2D *in vitro* studies have demonstrated that varying matrix stiffness changes the extent of branch initiation within mammary explants in engineered collagen I matrices^17^. Additionally, branch elongation and cell migration are guided by tensile forces that drive local fiber alignment^18,19^. The ECM protein fibronectin is critical in driving cleft formation in epithelial branching *in situ*^20^, and local accumulation of type I collagen at branch flanks and clefts regulates mammary branching *ex vivo*^21^. This demonstrates how ECM dynamics, both *ex vivo* and *in situ*, appear to modulate local mechanics similarly to what has been previously observed *in vitro*^17–19^. Collectively, these studies highlight the critical role of ECM mechanics in mammary branching.

While it is known that matrix stiffness regulates branching morphogenesis^22–24^, the effect of ECM viscoelasticity remains an open question. Biological tissues are viscoelastic^25,26^, meaning they exhibit both elastic solid-like behavior and viscous fluid-like behavior. In response to a constant deformation, viscoelastic materials will relax stress over time, while for purely elastic materials, stress will remain constant over time. ECM viscoelasticity has been shown to influence cell spreading^27,28^, proliferation^28,29^, migration^30,31^, and differentiation^32–34^. Very recently, ECM viscoelasticity has been shown to influence intestinal crypt morphogenesis, breast epithelial cell growth, and nephrogenesis in kidney organoids^35–37^. However, it remains unknown whether matrix viscoelasticity influences branching morphogenesis generally, including within mammary tissue, and the mechanisms underlying such processes are yet to be determined.

Here, we report the results of a systematic investigation into the role of matrix stress relaxation in regulating branching of human mammary epithelium. We explored biophysical mechanisms underlying mammary branch development and identified signaling pathways that are differentially regulated via matrix stress relaxation rate. This work reveals how matrix viscoelasticity regulates mammary branching, which can be leveraged for applications in tissue engineering and regenerative medicine, and may be relevant in other branching morphogenesis contexts as well.

## Results

### Slower matrix stress relaxation enhances mammary branching

We used tunable viscoelastic matrices to probe the impact of different stress relaxation rates on mammary branching. We formed 3D matrices using interpenetrating network (IPN) hydrogels composed of alginate and collagen I. Alginate is a bioinert polysaccharide that enables viscoelasticity to be tuned independently of stiffness by varying its molecular weight^38^. Divalent cations, such as calcium, form ionic crosslinks between alginate chains, which can break under stress and reform, allowing local matrix flow and giving rise to the stress relaxing properties of the hydrogel^39^ (Fig. 1A). Collagen I was incorporated as it is the major structural protein within the mammary gland^15^ and has been previously shown to induce elongation in mammary epithelial cells^40^. The stress relaxing properties of healthy mammary gland tissue are not well-characterized, so we fabricated hydrogels within a range of stress relaxation half times where cells are mechanically responsive^26^ (*t*_1/2_ ≈ 100 s, 800 s, 1200 s, 4000 s) (Fig. 1B) by using alginates of different molecular weights. The calcium concentration for alginate crosslinking was optimized to achieve a constant elastic modulus of approximately 200 Pa across all stress relaxation groups (Fig. 1C), mimicking physiological mammary tissue^41^. Importantly, we were able to tune the mechanical properties of the hydrogel independently of the collagen microarchitecture of the substrates (Fig. 1D).

**Figure 1:**
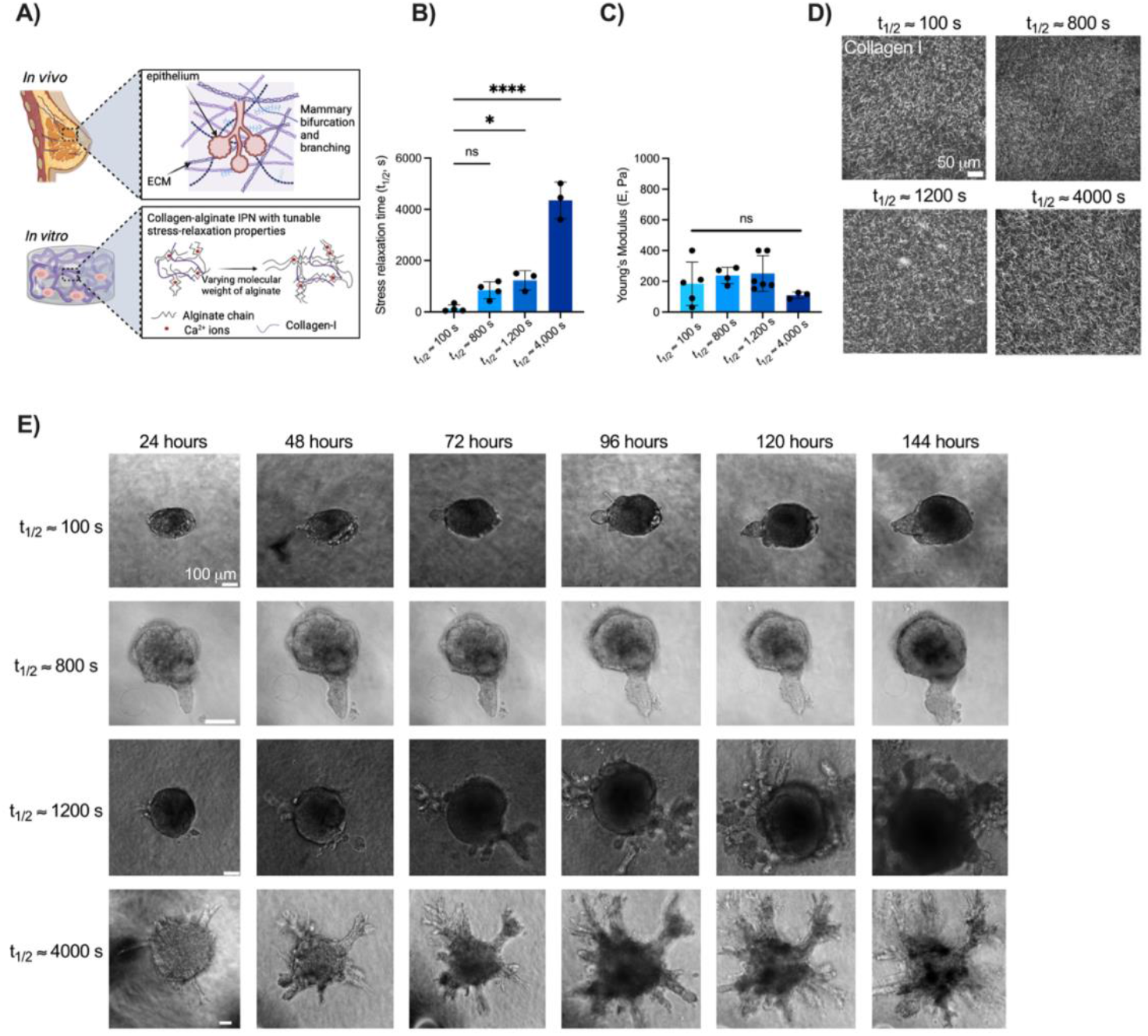
Matrix viscoelasticity regulates mammary branching of human epithelial spheroids. (a) Schematic depicting how mammary gland development is influenced by mechanical and biochemical cues within the *in vivo* microenvironment. 3D collagen-alginate interpenetrating networks (IPNs) enable independent tuning of the viscoelasticity of the matrix *in vitro*. Image created in *BioRender*. (b) Alginates of different molecular weights can be used to fabricate matrices with varying stress relaxation times. (c) Time sweep revealed that hydrogels with varying stress relaxation times had Young’s moduli that were not significantly different. (d) Representative confocal reflectance microscopy images depict collagen microarchitectures of matrices with varying stress relaxation times. (e) Brightfield images of MCF10A spheroids in 3D slow stress relaxing matrices (t_1/2_ ≈ 1200 s, t_1/2_ ≈ 4000 s) and fast stress relaxing matrices (t_1/2_ ≈ 100 s, t_1/2_ = 800 s) over 6 days. All data are represented as mean +/- standard deviation (SD). Statistical significance was determined using one-way ANOVA with Dunnett’s multiple comparison tests. Significance is indicated as follows: ****p < .0001, *p < .05, and n.s. = not significant.

We next assessed the influence of matrix stress relaxation on mammary branching. MCF10A human mammary epithelial cells, a cell line commonly used to model mammary development^42^, were formed into spheroids of approximately 3000 cells. MCF10A spheroids were encapsulated in alginate-collagen matrices with various stress relaxation half times (*t*_1/2_ ≈ 100 s, *t*_1/2_ ≈ 800 s, *t*_1/2_ ≈ 1200 s, and *t*_1/2_ ≈ 4000 s). Over the course of six days, tissues in fast stress relaxing conditions (*t*_1/2_ ≈ 100 s, 800 s) expanded isotropically and formed buds into the surrounding matrix (Fig. 1E). However, spheroids in slower stress relaxing conditions (*t*_1/2_ ≈ 1200 s, 4000 s) broke symmetry and branched extensively (Fig. 1E). Using confocal microscopy and fluorescently labeled cells, we observed compact and symmetric geometries of spheroids in fast stress relaxing matrices, but several branched structures stemming from the spheroid body in the slow stress relaxing matrices (Fig. 2A). This was characterized by a significant increase (586%) in the total number of branches (Fig. 2B) and a significant increase in branch length (390%) within slow stress relaxing matrices (*t*_1/2_ ≈ 1200 s) compared to fast stress relaxing matrices (*t*_1/2_ ≈ 100 s) after six days in culture (Fig. 2C). To account for experimental heterogeneity, we quantified the percentage of spheroids branching in any given experiment and found that spheroids in slow stress relaxing matrices exhibited the greatest branch frequency when compared to spheroids in fast stress relaxing matrices (Fig. 2D). Additionally, the cross-sectional area of the spheroid was not significantly different between stress relaxation groups after 24 hours in culture (Fig. 2E) but there was a significant increase in cross-sectional area (57%) after 120 hours in slow stress relaxing matrices (*t*_1/2_ ≈ 1200 s) (Fig. 2F). Spheroids in slow stress relaxing matrices were also significantly less circular than those in fast stress relaxing matrices, both at 24 and 120 hours (Fig. 2G, H). Our results demonstrate that matrix stress relaxation plays a key role in regulating mammary epithelial spheroid morphology and branch formation.

**Figure 2:**
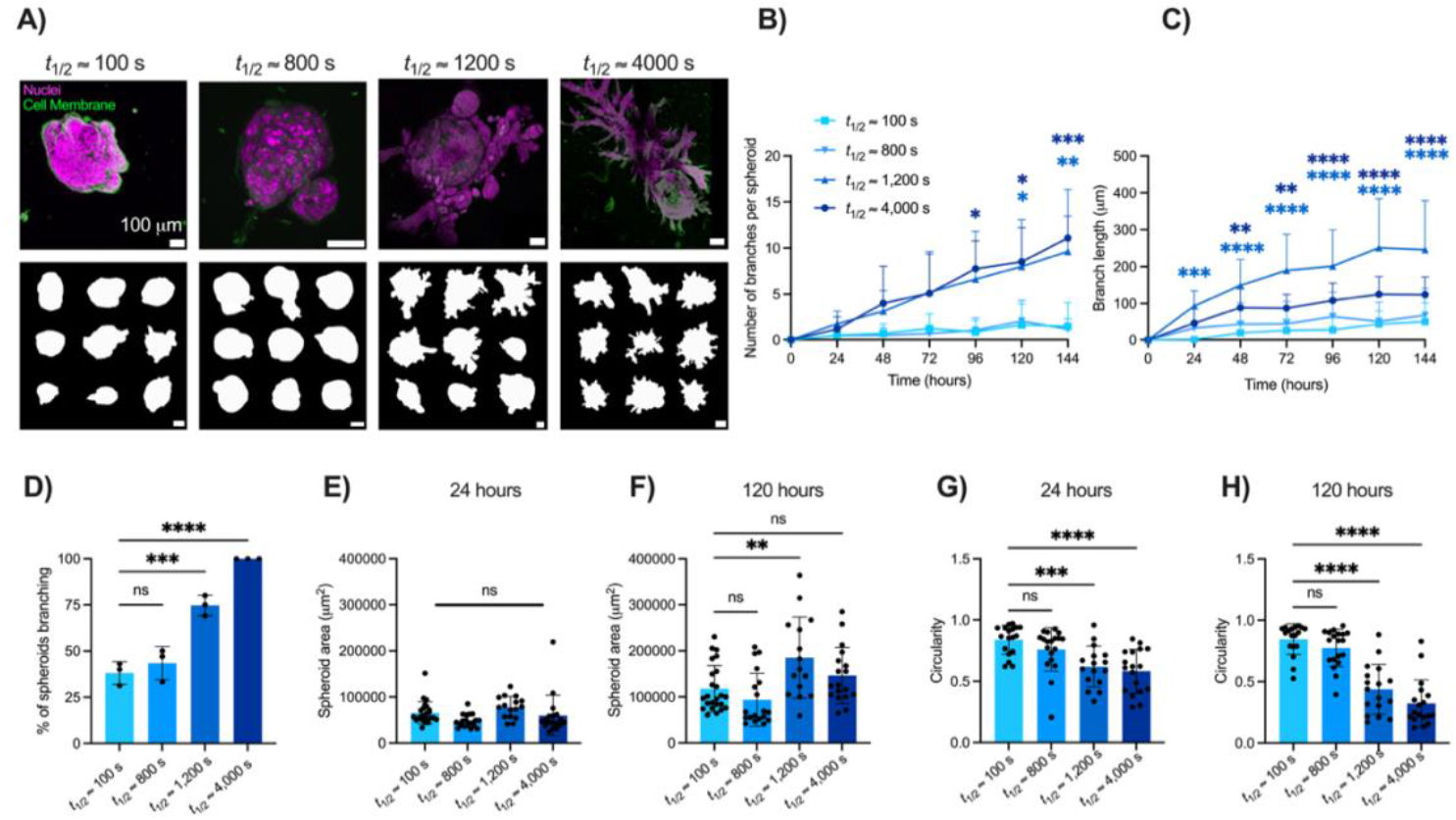
Mammary epithelial spheroid morphology and branch formation varies with stress relaxation rate. (a) Confocal images of a MCF10A spheroid in slow stress relaxing matrices (t_1/2_ ≈ 1200 s, t_1/2_ ≈ 4000 s) and fast stress relaxing matrices (t_1/2_ ≈ 100 s, t_1/2_ = 800 s). Cell membranes were labeled with rhodamine-B (green) and the nuclei with Hoechst 33342 (magenta). Representative outlines of MCF10A spheroid morphologies are portrayed after 120 hours in culture in varying stress relaxing matrices. (b) Quantification of branches revealed more branches were produced in slow stress relaxing matrices (t_1/2_ ≈ 1200 s, t_1/2_ ≈ 4000 s) and (c) this was accompanied by greater branch lengths. (d) Branch frequency was significantly enhanced in slow stress relaxing matrices (t_1/2_ ≈ 1200 s, t_1/2_ ≈ 4000 s), where each data point represents the percentage of spheroids that branched in an independent experiment, and quantified on day 7. (e) Quantification of the cross-sectional area of MCF10A spheroids at 24 hours and (f) 120 hours. (g) Quantification of circularity at 24 hours and (h) 120 hours in slow and fast stress relaxing matrices. Statistical significance was determined using one-way ANOVA with Šídák’s or Dunnett’s multiple comparison tests: ****p < .0001, p*** < .001, p** < .01, *p < .05, and n.s. = not significant, and performed with respect to the fast stress relaxing condition (t_1/2_ ≈ 100 s). n = 5-15 images from 3 hydrogels per condition. All data are represented as mean +/- SD.

### Slow stress relaxing matrices promote collagen fiber alignment along branching axis

We next sought to elucidate the biophysical mechanisms through which epithelium can generate forces to elongate branches and pattern epithelial trees. Previous work has found that collagen can accumulate around the outer flanks of mammary branches and at branch bifurcation sites^21^. Further, mammary gland explants will branch along a pre-aligned collagen network, following the direction of the collagen orientation axis^43^. To determine if viscoelasticity can drive mammary morphogenesis via changes in collagen fiber alignment, we employed confocal reflectance microscopy to visualize the collagen network surrounding epithelial branches in both fast (*t*_1/2_ ≈ 100 s) and slow stress relaxing matrices (*t*_1/2_ ≈ 1200 s, 4000 s). We then calculated the orientation angle (0°–90°) between the spheroid boundary and local collagen fibers. Regions of interest were drawn to delineate collagen fiber alignment along the branching axis and adjacent to the branching axis^40^. Confocal reflectance images and a density heat map demonstrated enhanced collagen fiber alignment in slow stress relaxing matrices along the branching axis (Fig. 3A). Upon binning and normalizing the orientation values, we observed significantly increased collagen fiber alignment (corresponding to bins between 60–90*°*) in front of the branching axis in slow stress relaxing matrices, but not in fast stress relaxing matrices (Fig. 3B–D). The collagen fiber orientation values adjacent to the mammary branches (away from the tip) were uniformly distributed regardless of matrix stress relaxation (Supplementary Fig. 1A–C), indicating a random collagen fiber network. Thus, our findings suggest that mammary epithelial cells spatially remodel their collagen matrix to promote branching morphogenesis in a stress relaxation-dependent manner.

**Figure 3:**
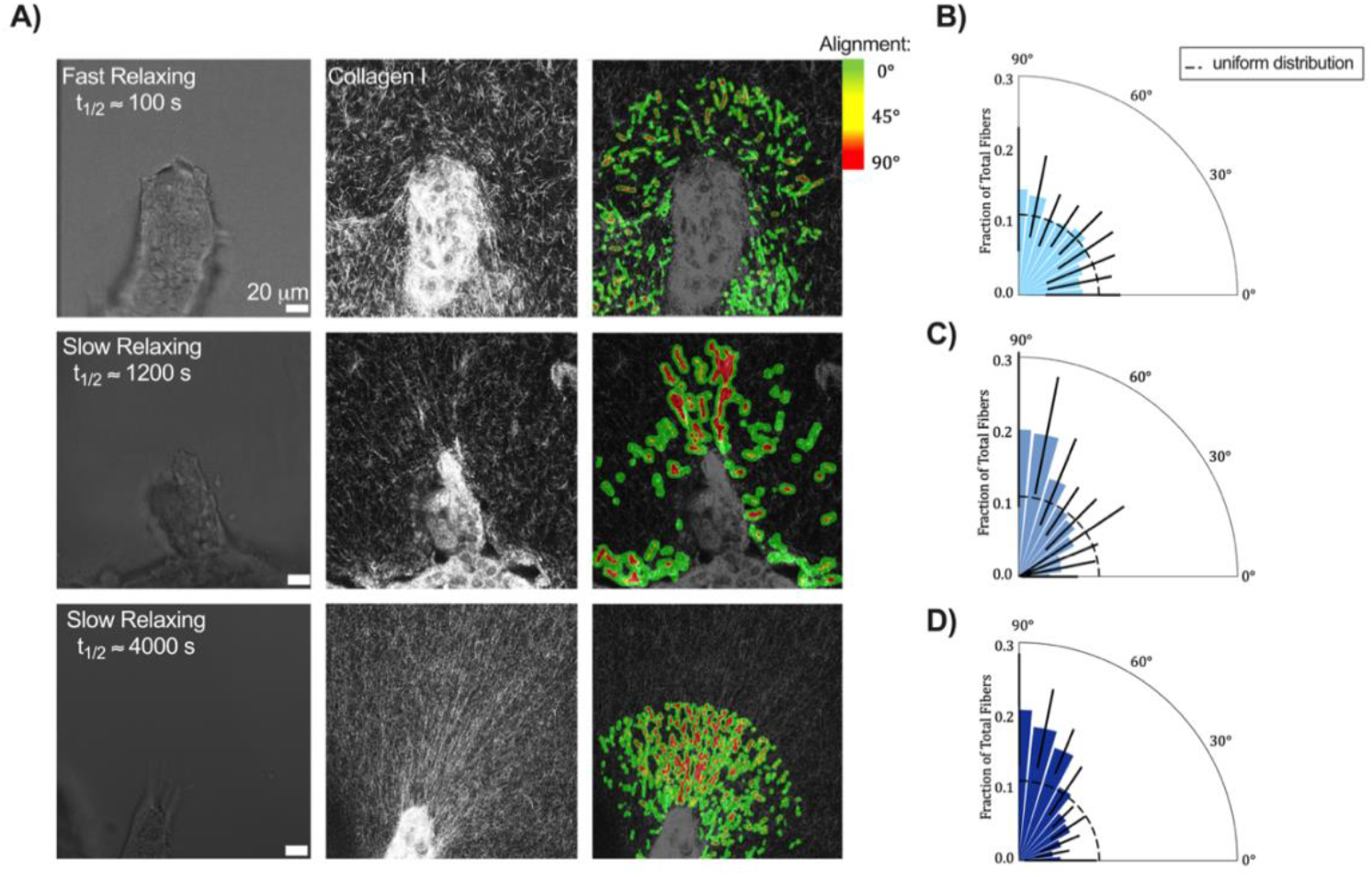
Slow relaxing matrices promote collagen fiber alignment along branching axis. (a) Representative brightfield and confocal reflectance images of the collagen fiber network surrounding a MCF10A spheroid branch in various stress relaxing conditions (t_1/2_ ≈ 100 s, t_1/2_ ≈ 1200 s, t_1/2_ ≈ 4000 s). Heatmap depicts the extent of collagen fiber alignment via *CurveAlign*. (b) Quantification of the relative orientation angle along the branching axis demonstrated that collagen fibers are randomly aligned in the t_1/2_ ≈ 100 s gel, (c) highly aligned in the t_1/2_ ≈ 1200 s gel, and (d) also highly aligned in the t_1/2_ ≈ 4000 s gel. The dashed line represents the expected value if all fibers were uniformly distributed within the bins. n = 9 images from 3 hydrogels per condition. Black error bars indicate SD for each group. Statistical analysis conducted using one-way ANOVA with a Kruskall-Wallis test to compare the fraction of fibers in each bin in C and D with the corresponding bin in B. **p < .01 between all groups with angles greater than 60 degrees.

### Cell-generated forces displace the matrix to facilitate mammary branching in slow stress relaxing matrices

Since we observed enhanced collagen fiber alignment in front of the branches in slow stress relaxing matrices, we next evaluated the extent of matrix deformation during branching. We encapsulated MCF10A spheroids in both slow (*t*_1/2_ ≈ 1200 s) and fast (*t*_1/2_ ≈ 100 s) stress relaxing conditions along with 0.2 μm fluorescent beads for 48 hours and tracked the bead displacement over the next 16 hours in culture (Fig. 4A, B). Spheroids in fast stress relaxing conditions generated small, isotropic pushing displacements in the surrounding matrix (Fig. 4A, Supplementary Video 1). Interestingly, branch tips in slow stress relaxing matrices generated displacements in the pulling direction during branch extension, whereas branch bifurcation sites exhibited pushing displacements (Fig. 4B, Supplementary Video 2).

**Figure 4:**
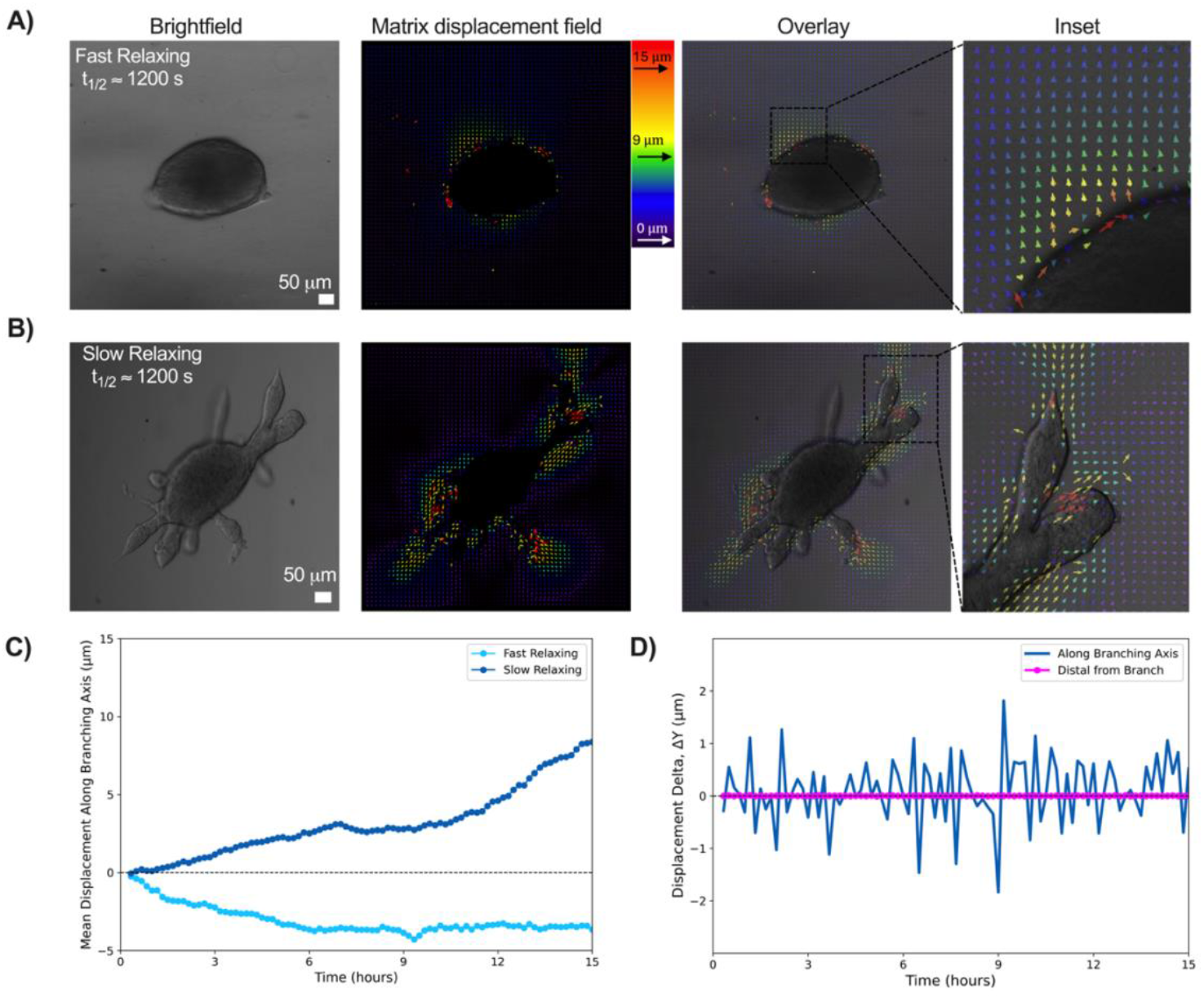
Cell-generated non-continuous contractions displace the matrix to facilitate mammary branching in slow stress relaxing matrices. (a) Matrix displacement field of a MCF10A spheroid in a fast stress relaxing matrix demonstrated minimal displacements were applied. (b) Matrix displacement field of a MCF10A spheroid in a slow stress relaxing matrix demonstrated high matrix displacements were applied in front of branches. Matrix displacement field generated via particle image velocimetry (PIV) Image J plugin. (c) Positive cumulative mean displacements of greater magnitude were observed in slow stress relaxing matrices suggesting pulling forces, while fast stress relaxing matrices showed smaller, negative displacements, suggesting pushing forces. (d) Quantification of mean displacement delta demonstrated mammary epithelium exert non-continuous contractions to extend their mammary branches in slow stress relaxing matrices. Bead displacement position data acquired from *Imaris* and analyzed and plotted in MATLAB.

This demonstrates how cells spatially pattern the application of force to the ECM to enable branch extension and bifurcation. We next quantified the direction and magnitude of displacements with respect to a reference frame where the y-axis was oriented toward the branch, so that any displacements applied directionally toward the branch would result in a positive y-axis displacement, and any displacements away from the branch would result in negative y-axis displacement. Interestingly, we found that larger cumulative displacements occurred in front of branch tips in slow stress relaxing matrices, compared to their fast stress relaxing counterparts (Fig. 4C). We observed that the displacements were both positive and of greater magnitude in slow stress relaxing matrices than in fast stress relaxing matrices (Fig. 4C). This indicates that mammary epithelium exerts pulling forces (as demonstrated by the positive displacement field) to extend branches in slow stress relaxing matrices, but in fast stress relaxing matrices, spheroids exert pushing forces (as demonstrated by the negative displacement field). Collectively, these results demonstrate how the cell-generated forces that drive tissue growth and branching are dependent on the stress relaxation of the matrix. We next looked at the changes in displacement between consecutive frames (delta displacement) in front of a representative branch within a slow stress relaxing matrix. We found an intermittent increase and decrease over time, compared to a region of interest distal from the branch, where there was no change in delta displacement over time (Fig. 4D). This illustrates that cells exert pulling forces in an intermittent, non-continuous manner in slow stress relaxing matrices, and these forces are applied in front of elongating branches.

### Actomyosin contractility and phosphorylated focal adhesion kinase contribute to mammary branch elongation

Given that we observed enhanced collagen fiber alignment at the leading edge of elongating mammary branches, and that mechanical forces are sufficient to orient and align collagen fibers^44^, we hypothesized that cell contractility drives collagen fiber alignment, and thereby mammary epithelial branch elongation. We investigated the role of β1 integrin, focal adhesion kinase activity, and actomyosin-based contractility on mammary branching, as these are key components of mechanoresponsive pathways that enable cellular force generation. Integrin binding could enable force transmission to align collagen fibers, and this would be associated with downstream signaling proteins, including focal adhesion kinase (FAK)^45^. In particular, we examined β1 integrins, as they are known to play a role in collagen fiber bundling^46^ and are implicated in mammary branching^47^. An examination of the localization of β1 integrin and phosphorylated FAK (Tyr397) via immunostaining and confocal microscopy revealed significant increases (74% and 53%, respectively) in the abundance of these proteins at the tip cells of mammary branches in slow stress relaxing matrices compared to the abundance in the spheroid body (Fig. 5A–C).

**Figure 5:**
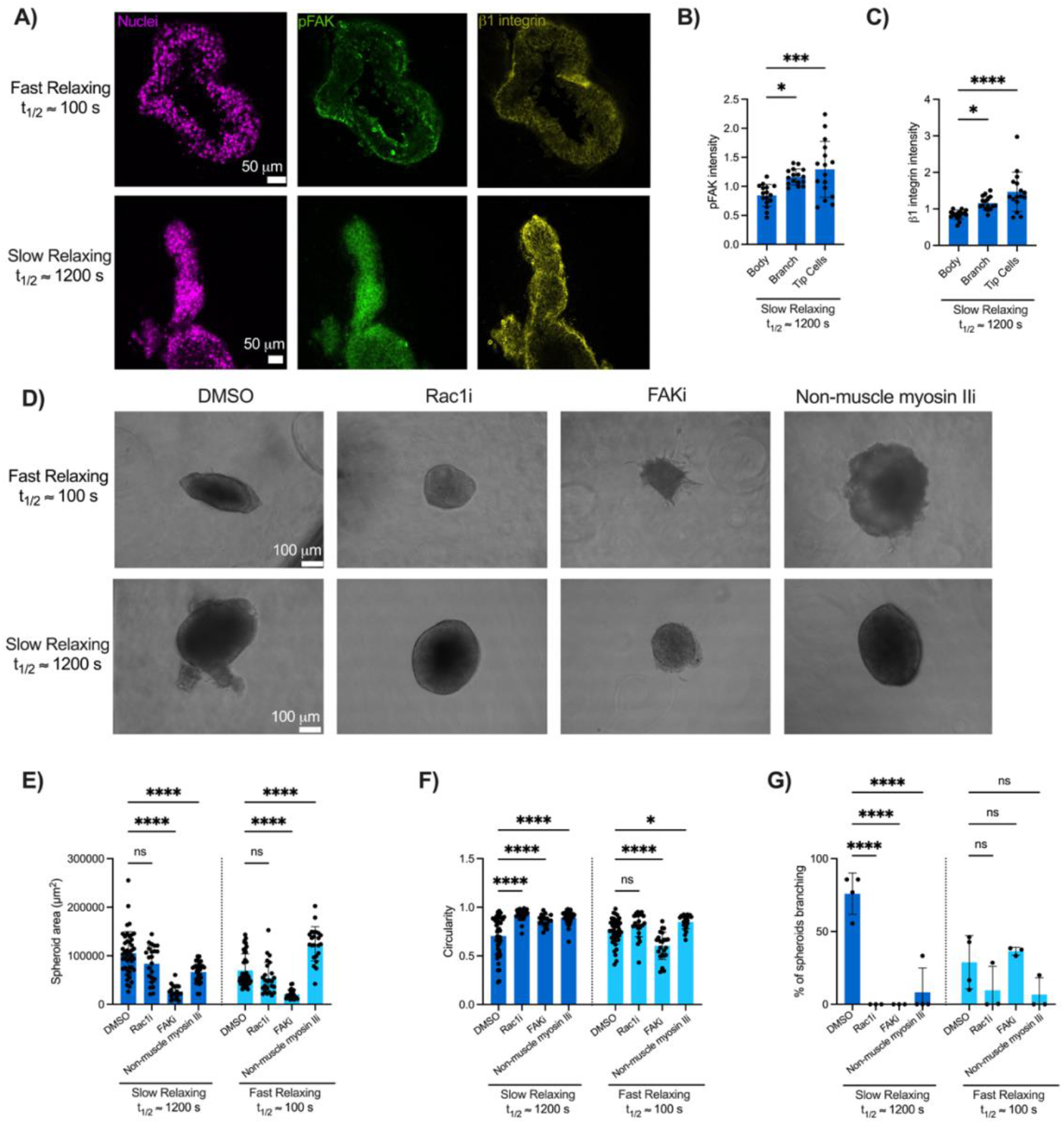
Actomyosin contractility and phosphorylated focal adhesion kinase contribute to mammary branch elongation. (a) Immunohistochemical stains of phosphorylated FAK (Tyr397) and β1 integrin in MCF10A spheroids in both fast (t_1/2_ ≈ 100 s) and slow (t_1/2_ ≈ 1200 s) and stress relaxing matrices after 7 days in culture. (b) There was greater abundance of both FAK phosphorylation (pFAK) and (c) β1 integrin at the tip cells of a mammary duct compared to the spheroid body in slow relaxing gels. Image intensity values were normalized to the nuclei stain (Hoechst 33342). (d) Representative brightfield images of MCF10A spheroids cultured in fast and slow stress relaxing gels after 7 days with the treatment of Rac1, FAK, or non-muscle myosin II inhibitors, or DMSO as a vehicle control. (e) Quantification of spheroid area, (f) circularity, and (g) branch frequency when cultured in indicated conditions. Statistical significance was determined using one-way ANOVA and with Tukey’s or Šídák’s multiple comparison tests: ****p < .0001, ***<p.001, **p<.01, *p<.05, and n.s. = not significant. n = 5-15 images from 3-5 hydrogels per condition. All data are represented as mean +/- SD.

Next, we investigated the hypothesis that the ability of mammary epithelial cells to undergo branching morphogenesis is dependent on cell contractility and force-generating pathways. Non-muscle myosin II is a motor protein that enables cell contractility and mechanical stability, and Rac1, a GTPase that regulates the formation and maintenance of focal adhesions, has been shown to regulate branch elongation and tissue growth^36,48^. We pharmacologically inhibited FAK, non-muscle myosin II, and Rac1, and found that inhibition of FAK led to a significant decrease in spheroid area in both slow and fast stress relaxing matrices (75% and 70%, respectively) compared to untreated controls (Fig. 5D, E). Inhibition of non-muscle myosin II led to a significant decrease (37%) in spheroid area in slow stress relaxing matrices compared to untreated controls, but a significant increase (78%) in spheroid area in fast stress relaxing matrices (Fig. 5E). Interestingly, inhibiting Rac1 resulted in no significant effect on spheroid area in slow or fast stress relaxing matrices (Fig. 5E). In slow stress relaxing matrices, circularity was significantly increased when Rac1, FAK, and non-muscle myosin II were inhibited (31%, 22%, and 27%, respectively), although in fast stress relaxing matrices, inhibiting FAK led to a 20% significant decrease in circularity due to the formation of cellular protrusions throughout the spheroid (Fig. 5F). Further, inhibition of all three signaling pathways led to a significant decrease in branching frequency in slow stress relaxing matrices, but there was no significant difference within fast stress relaxing matrices (Fig. 5G). Collectively, these results show that β1 integrin, phosphorylated FAK, Rac1, and non-muscle myosin II play key roles in contributing to mammary morphogenesis in slow stress relaxing matrices.

### Hyperosmotic stress reduces mammary spheroid growth and branching in slow stress relaxing matrices

ECM viscoelasticity relates strongly to the concept of confinement^49^, as it dictates the extent to which cells can deform and remodel their microenvironment over time. Thus, we next probed the role of confinement on mammary branching by applying hyperosmotic stress to mammary epithelial spheroids. Previous work has shown that the pressure of fluid within the developing lung can drive morphogenesis^50^ and breast epithelial cellular membrane breaching is driven by cell volume expansion^51^. Extracellular matrix mechanics impact single cell volume expansion, as recent work has found that A7 cells will decrease their water content in response to an increased substrate stiffness^52^, and mesenchymal stem cell (MSC) volume expansion is regulated by stress relaxation^34^. Here, we applied hyperosmotic stress to spheroids by the addition of varying concentrations of 400 Da polyethylene glycol (PEG) in growth media, as in prior studies^52,53^. After a week of culture under hyperosmotic pressure, we found that mammary branching was significantly impeded in both slow and fast stress relaxing matrices compared to unperturbed controls (Fig. 6A). We observed that both low (Δ *P* = 92 kPa, 37.5 mOsm/L) and high osmotic pressures (Δ *P* = 197 kPa, 75 mOsm/L) led to a significant decrease in cross-sectional spheroid area in both slow and fast stress relaxing matrices compared to the control (Δ *P* = 0 kPa, 0 mOsm/L) (Fig. 6B). Additionally, low (Δ *P* = 92 kPa) and high osmotic pressures (Δ *P* = 197 kPa) significantly enhanced spheroid circularity in slow stress relaxing matrices (Fig. 6C). Only low osmotic pressure (Δ *P* = 92 kPa) enhanced circularity in fast stress relaxing matrices, although spheroids in fast stress relaxing matrices were already very circular (Fig. 6C). Furthermore, branch length and branch number were significantly reduced in slow stress relaxing matrices under both low (Δ *P* = 92 kPa) and high (Δ *P* = 197 kPa) osmotic pressures (Fig. 6D, E). Interestingly, we found that applying varying levels of hypoosmotic stress (Δ *P* = −155 kPa, Δ *P* = −310 kPa) for a week in culture was not sufficient to enhance mammary growth or branching in either slow or fast stress relaxing matrices (Supplementary Fig. 2). To further investigate the extent to which hyperosmotic pressure impairs mammary branching, we dynamically modulated the applied pressure within our cell culture system. Spheroids were first allowed to grow for four days in culture under normal conditions, after which hyperosmotic pressure was applied and maintained for the following three days. We found that applying low osmotic pressure (Δ *P* = 92 kPa) after four days in culture impaired the ability for spheroids to grow and form branches in slow stress relaxing matrices for the subsequent three days (Fig. 6F–H, Supplementary Fig. 3A). This trend was also consistent for spheroids cultured in fast stress relaxing matrices (Fig. 6I–K, Supplementary Fig. 3B). This effect was further amplified when we applied higher osmotic pressures (Δ *P* = 197 kPa) after four days in culture, and we found that higher osmotic pressures were sufficient to halt both branching and cellular expansion in both slow and fast stress relaxing matrices (Supplementary Fig. 4). Additionally, we conducted a complementary experiment where osmotic pressure was applied for the first four days in culture, followed by a return to normal conditions for the remaining three days. We found that when the lower osmotic pressure (Δ *P* = 92 kPa) was released after four days in culture, spheroids were able to grow but failed to resume branching in both slow and fast stress relaxing matrices (Supplementary Fig. 5A–H). When we performed the same experiment with higher osmotic pressures (Δ *P* = 197 kPa), we found that spheroids were unable to recover branching in both slow and fast stress relaxing matrices after osmotic pressure was released on day four (Supplementary Fig. 5I–P). In summary, our findings demonstrate that changes in osmotic pressure can disrupt mammary morphogenesis, even after this pressure is subsequently released.

**Figure 6:**
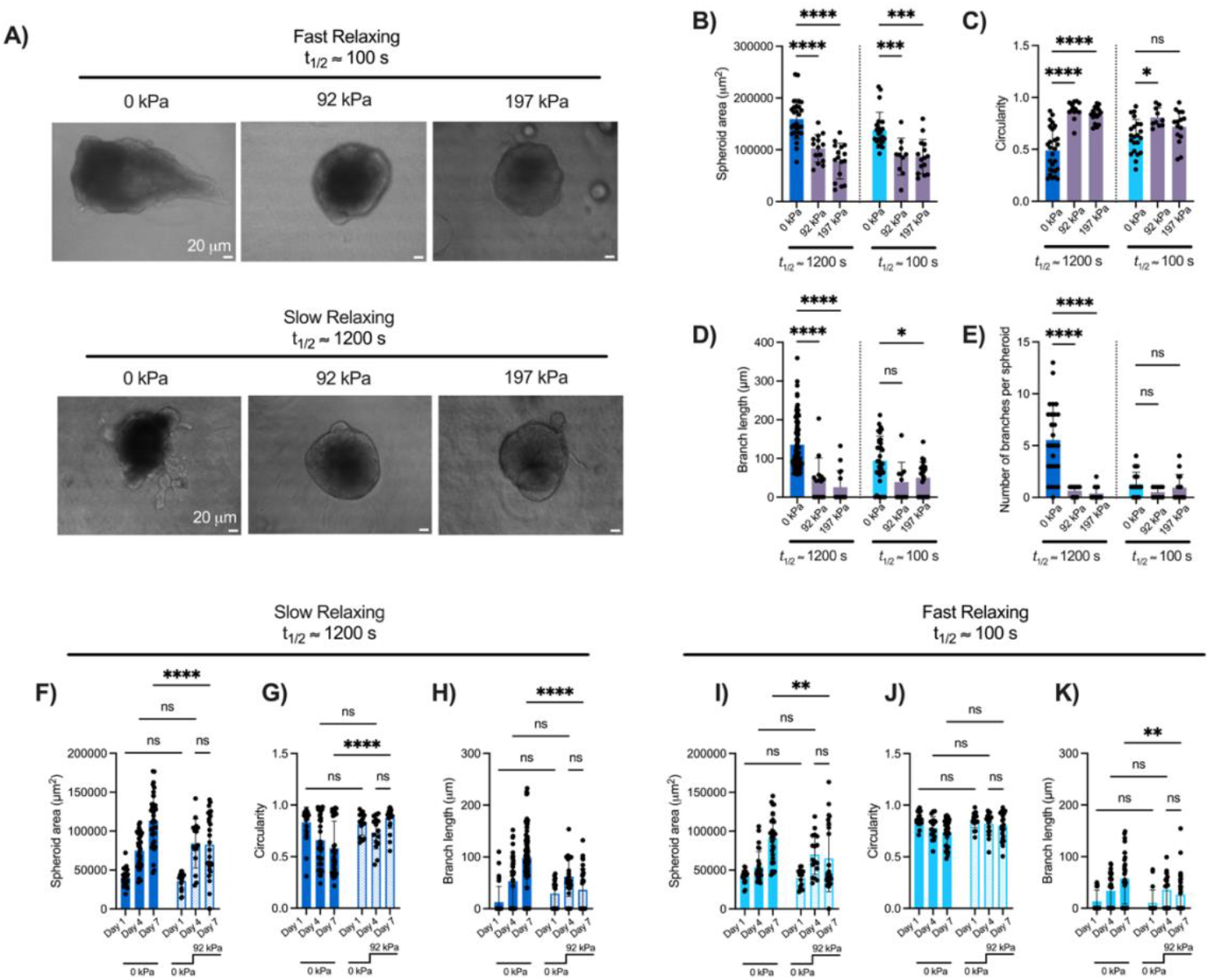
Hyperosmotic stress modulates mammary spheroid growth and branching in slow stress relaxing matrices. (a) Brightfield images of MCF10A spheroids in various stress relaxing conditions under continuous hyperosmotic stress for 7 days. (b) Quantification of MCF10A spheroid area under varied osmotic pressure (Δ *P* = 0, 92, 197 kPa) for 7 days, in both slow (t_1/2_ ≈ 1200 s) and fast (t_1/2_ ≈ 100 s) relaxing matrices. (c) Circularity was enhanced in slow stress relaxing gels when spheroids were subjected to hypertonic media. (d) Quantification of branch length demonstrated that spheroids branched to a lesser extent in slow stress relaxing gels under an osmotic pressure of Δ *P* = 92 and 197 kPa. (e) In slow stress relaxing gels, spheroids branched less when subjected to hyperosmotic pressure, but there was no significant difference in fast stress relaxing gels. (f) MCF10A spheroid growth in slow stress relaxing matrices was halted when spheroids were subjected to hyperosmotic pressure on day 4 in culture, and this was accompanied by (g) increased circularity and (h) decreased branch length. (h) MCF10A spheroid growth in fast stress relaxing matrices was halted when spheroids were subjected to hyperosmotic pressure on day 4 in culture. (i) There was no significant difference in circularity when osmotic pressure was applied on day 4 in culture in fast stress relaxing matrices, although (k) branch length decreased significantly compared to the control. Statistical significance was determined using one-way ANOVA with Šídák’s multiple comparison tests: ****p < .0001, p*** < .001, p**<.01,*p<.05 and n.s. = not significant. n = 5-15 images from 3 independent experiments. All data are represented as mean +/- SD.

This suggests that transient increases in confinement, whether through osmotic pressure or changes in ECM viscoelasticity, play a critical role in regulating mammary epithelial branching.

## Discussion

Here, we provide evidence that matrix stress relaxation is a key parameter in regulating mammary branching morphogenesis. Our unique approach allows us to decouple matrix stress relaxation from matrix stiffness and collagen fiber architecture to better understand how stress relaxation independently governs mammary branching. Our study reveals that cell-generated forces, driven by actomyosin contractility and focal adhesion kinase activity, facilitate branch elongation in slow stress relaxing matrices. Our results suggest that mammary epithelium locally remodel the collagen fiber network and displace the ECM to enable branch elongation. Importantly, we find that this behavior only emerges in slow stress relaxing matrices, thereby demonstrating the importance of the timescale of stress relaxation in tissue morphogenesis.

While the influence of matrix stiffness on tissue morphogenesis has been increasingly recognized^22–24^, the mechanism of how viscoelasticity mediates morphogenesis remains poorly understood. Here, we show that branching morphogenesis is supported in slow stress relaxing matrices, but not in fast stress relaxing matrices. Although the stress relaxing properties of healthy mammary gland tissue are not well-characterized, the mammary gland is surrounded by adipose tissue, which exhibits relatively slow stress relaxation, on the order of hundreds of seconds^26^. This may, in part, explain why the slow stress relaxing matrices promote mammary branching. Prior studies have found that cells migrate collectively via local proteolysis of the matrix or ECM softening^19,54^, and our findings raise the possibility that mechanical confinement in slow stress relaxing matrices may spatially constrain cells, thereby promoting branch initiation in local regions of heterogeneity. A recent report found that mammary epithelial spheroids undergo symmetry breaking and invade the surrounding matrix when embedded in stiff (5,000 Pa) and fast stress relaxing alginate matrices^36^. We hypothesize that the differences in MCF10A spheroid behavior are attributable to two key distinctions in the microenvironmental properties between these studies; i) the matrices used in our experiments were over an order of magnitude softer (200 Pa vs. 5,000 Pa), and ii) we incorporated fibrillar collagen I within the matrix for cell adhesion, instead of a direct conjugation of the RGD adhesion motif to the alginate. These distinctions highlight the critical role of matrix composition and matrix stiffness in regulating cell behavior and morphogenetic processes.

While the role of the extracellular matrix in guiding epithelial form and function has long been recognized, the specific role of collagen I in directing mammary branch extension has been contested. In murine models, prior studies have demonstrated that patterning of collagen fibers in the mammary gland stroma regulates epithelial orientation during development^43^ and that cell-induced matrix alignment facilitates collective cellular migration^55^. It has been proposed that ECM fibers may serve as conduits for long-range stress transmission, thereby enabling mechanotransduction across tissue scales^56^. Conversely, recent reports suggest that directional epithelial outgrowth occurs independently of collagen I alignment^21^, and computational simulations suggest branching can occur in the absence of long-range guidance cues from the ECM^57^. Here, we show that human mammary epithelia apply directional forces to mechanically constrain and stabilize the ECM and guide branch development, which is consistent with recent studies that show that collagen fiber alignment is generated by expanding branches of human mammary organoids^18^. While the temporal sequence of these events remains to be fully resolved, our findings support a model of mechanical feedback between epithelial dynamics and ECM organization which is mediated by the viscoelastic properties of the matrix.

Notably, we have also shown in this study how integrin-mediated, non-continuous displacements are applied to the ECM to drive branch elongation. Previous work has suggested the importance of phosphorylated FAK and integrin activity in regulating collagen fiber alignment in stress relaxing matrices, particularly within the context of human MSCs^27^. Additionally, β1 integrin signaling is required for directional migration during mammary morphogenesis^58^. Our work builds upon this foundation by showing that both FAK and Rac1 are critical for mammary branching in slow stress relaxing matrices, and that phosphorylated FAK and *β*1 integrin localize to the tip cells of extending branches. Given the role of integrins in coupling ECM adhesions to the actin cytoskeleton, these findings suggest a mechanism by which localized signaling at the branch tips enables cells to generate and transmit contractile forces to the surrounding matrix. Indeed, the role of contractile and tensile forces has been extensively studied in tissue development and morphogenesis^59^, and our observations support a model in which actomyosin-dependent contractions contribute to tissue extension^19,60^, which occurs in a non-continuous manner^18^. This dynamic behavior is reminiscent of pulsed actin assembly during collective cell migration^61,62^, suggesting that temporally regulated force generation may be a conserved mechanism across diverse cellular and developmental processes.

We next evaluated how osmotic confinement regulates mammary branching and found that applying hyperosmotic pressure resulted in the inhibition of branch formation in slow stress relaxing matrices. As hyperosmotic stress reduces intracellular pressure, our work aligns with other studies that have found that morphogenetic processes rely on hydrostatic pressure to drive cellular expansion^63^. *In vivo*, confinement can arise from changes in basement membrane composition, stiffness, or mechanical stability, and is known to play a critical role in shaping tissues by influencing cell behavior, tissue organization, and the coordination of intrinsic mechanical cues^11^. In the context of mammary branching, it is known that the geometry and confinement of growing mammary tissues dictates their branching pattern^18,22^. Within broader epithelial morphogenetic processes, changes in osmotic gradients are correlated to lumenogenesis and epithelial cell proliferation^64,65^ and cells generate intracellular tension to drive the elongation of branches in the developing mouse lung^66^. Although the application of hypoosmotic pressure did not enhance mammary branching, it is still possible that mammary branching is driven by changes in internal pressure, such as cell proliferation or other osmotic fluctuations. Hypoosmotic pressure was induced using previously established methods^34,51^ where the growth medium was diluted with DI water. However, we acknowledge that this dilution could introduce confounding effects on the cells. Our findings contribute to a growing body of literature that cell volume regulation plays a key role in cellular processes such as mammary branching. Recent studies have shown that MSC volume expansion during cell spreading activates TRPV4 ion channels to enhance osteogenic differentiation^34^, and cell volume regulation is driven by both chloride ion channels and ATP-dependent processes^52^. Further exploration of the role of TRP channels, mechanosensitive ion channels, and aquaporins in regulating and maintaining the pressure within mammary epithelium during branching could provide valuable insights into the mechanobiological processes underlying tissue morphogenesis. Prior to this work, the effect of matrix viscoelasticity on branching morphogenesis generally, and on mammary branching specifically, remained unknown. Here, we highlight that matrix viscoelasticity is critical in regulating mammary branching morphogenesis. Importantly, we elucidate that mammary branching is mechanistically driven by dynamic extracellular matrix remodeling and intermittent epithelial contractions that is regulated via focal adhesion kinase and β1 integrin signaling. Looking forward, this framework can be expanded upon to further understand how branching morphogenesis is regulated by mechanical cues in other glandular epithelia, such as the lungs and kidneys. More broadly, these insights have implications for tissue engineering and regenerative medicine, where tuning of matrix mechanics could be leveraged to direct morphogenetic outcomes.

## Materials and Methods

### A. Alginate Preparation

Sodium alginate with a high molecular weight (FMC Biopolymer, Protanal LF 20/40, 280 kDa) was used to fabricate the slow stress relaxing matrices (*t*_1/2_ ≈ 1200 s, 4000 s) and alginate with a low MW (UP VLVG, Pronova, 4200501 and UP LVG, Pronova, 4200001) was used to prepare fast stress relaxing matrices (*t*_1/2_ ≈ 100 s, 800 s, respectively). Alginates were dissolved in deionized water (1% w/v) and then dialyzed with tubing membrane (10 kDa MWCO) against deionized water for 3 days. The alginate solution was treated with activated charcoal, filtered (Sigma-Aldrich), sterile filtered (MilliporeSigma), frozen, lyophilized, and then dissolved in Dulbecco’s Modified Eagle Medium: Nutrient Mixture F–12 (DMEM/F–12, ThermoFisher).

### B. Cell Culture and Spheroid Formation

MCF10A cells (CRL-10317, ATCC) were cultured in Dulbecco’s Modified Eagle’s Medium/Nutrient Mixture F–12 (DMEM/F–12, ThermoFisher), and supplemented with 5% horse serum (Thermo Fisher), 20 ng/mL epidermal growth factor (Peprotech), 10 μg/mL insulin (Sigma), 0.5 μg/mL hydrocortisone (Sigma), 100 ng/ml cholera toxin (Sigma), and 1% penicillin/streptomycin (ThermoFisher), as previously described^67^. MCF10A cells were passaged at 70% confluency, typically every three to four days, and the media was changed every 48 hours. Cells were formed into MCF10A spheroids via the hanging-drop method^68^. Briefly, 0.68% w/v rat tail collagen I solution (Advanced Biomatrix) was added to growth medium and 15 μl droplets were pipetted onto the lid of a petri dish at a seeding density of 200 cells/μl. The petri dish was inverted, and 5 mL of phosphate-buffered saline (ThermoFisher) was added to the base of the petri dish to keep the spheroids hydrated. The petri dish was kept in a 5% CO_2_ humidified incubator at 37°C for 24–36 hours to allow for spheroid formation.

### C. Alginate-Collagen Hydrogel Fabrication and Spheroid Encapsulation

Hydrogels were fabricated using a syringe-mixing method that has previously been described^69^. First, warmed alginate was added into a 1 mL Luer lock syringe (Cole-Palmer), avoiding the creation of air bubbles^69^. Extra DMEM/F–12 (ThermoFisher) was added such that all matrices had a final alginate concentration of 0.5% w/v, except for the fabrication of the slow stress relaxing matrix (*t*_1/2_ ≈ 4000 s), which had a final concentration of 1.5% w/v. Next, calcium sulfate dihydrate (Sigma-Aldrich) was dissolved in deionized water, autoclaved, and subsequently added to a separate 1 mL Luer lock syringe (Cole-Palmer) such that the final concentration was 2 mM. Rat tail collagen I (Advanced Biomatrix) was kept on ice, and added to the syringe such that the final concentration was 2 mg/mL. The collagen was immediately neutralized with 10 mM sodium hydroxide (NaOH). The two syringes were then coupled with a Luer lock (Cole-Palmer), and rapidly mixed with 20 pumps on the syringe handles. For mechanical characterization tests, these substrates were deposited directly onto the parallel plate of the rheometer. For cell culture experiments, these gels were deposited directly into a sterile 48 well plate (Fisher Scientific) after spheroids were counted and added into the syringe with warmed alginate. The well plate was then immediately transferred to a 37°C incubator, and the gel was allowed to polymerize for one hour prior to adding cell-culture medium. Culture medium was changed every two days.

### D. Mechanical characterization of alginate-collagen IPNs

Mechanical properties of the hydrogel were assessed using *in situ* testing with an Anton Paar MCR 502 stress-controlled rheometer. Briefly, alginate hydrogels with low molecular weight alginate and high molecular weight alginate were fabricated, as described above. A 25 mm plate was lowered onto the polymer solution until the rheometer registered a non-negative axial force immediately after crosslinking commenced. Gels were maintained at 37 °C. A thin layer of mineral oil was then pipetted around the hydrogel to ensure the gels would not dehydrate during mechanical measurements. Once the storage modulus reached an equilibrium value, a stress relaxation test was performed. While the strain (10%) was held constant, stress was recorded over time. The stress relaxation time was defined as the time taken for the maximum stress to relax to half of its initial value. The storage and loss modulus of the hydrogel were measured using a time sweep test (1 Hz, 1% strain). From this data, the complex modulus (Equation 1) and elastic modulus were calculated (Equation 2). The complex shear modulus (G*) describes the material’s response to shear stress, whereas the elastic modulus (E) describes the material’s response to uniaxial stress. A Poisson ratio (*v*) was assumed to be 0.5^69^.

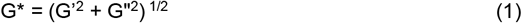

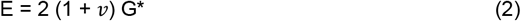

### E. Microscopy

All confocal images were collected using a laser scanning confocal microscope (Leica SP8). All brightfield images were collected using an Olympus IX50 – S8F2 inverted microscope. In live-cell time-lapse imaging, a Leica SP8 confocal microscope was utilized and equipped with an Okolab incubation chamber (37°C and 5% CO_2_). MCF10A spheroids were encapsulated with 0.2 μm dark red carboxylate-modified fluorescent microspheres (Molecular Probes) at a density of 1.6 x 10^12^ beads/mL and dispensed into chambered cover glasses (ThermoFisher). Spheroids were imaged every 10 minutes overnight using a Leica HC PL Fluotar 10x/0.30 objective.

### F. Immunofluorescence of fixed cells

To visualize the collagen network of the collagen-alginate IPNs, spheroids were first fixed on day four using 4% paraformaldehyde for 45 minutes at 37°C. The gels were then washed three times with Dulbecco’s phosphate-buffered saline with calcium (cPBS) for 20 minutes each time and then transferred to a glass cover slip. After this, a Leica 25x immersion objective was utilized to perform confocal reflectance microscopy. For the phosphorylated FAK and β1 integrin immunofluorescence analysis, all spheroids were fixed and washed as previously described. After this, they were left overnight in a 30% w/v sucrose in cPBS solution to dehydrate the gels. The following day, the gels were transferred to a solution of 50% w/v OCT and 50% w/v of the 30% w/v sucrose-cPBS solution for six hours to allow the OCT to penetrate the gel. Gels were then transferred into cryomolds with OCT and allowed to freeze on dry ice before being transferred to - 20°C. The samples were then sectioned with a cryostat (Leica) to 60 μm sections and transferred to glass Superfrost Plus slides (Thermofisher). The sections were then washed with cPBS and then incubated in a blocking buffer for one hour to minimize non-specific staining. Samples were then incubated with primary antibodies overnight, washed with cPBS, and then incubated with secondary antibodies for one hour. Primary antibodies used: β1 integrin *1:200* (Thermofisher, 14-0299-82), FAK-P-Y397 *1:500* (Thermofisher, 700255). Secondary antibodies used: Alexa Fluor Goat Anti-Ms IgG1 *1:1000*, 647 (Thermofisher, A21240), Alexa Fluor 488 Goat Anti-Rb IgG1 *1:1000* (Thermofisher, A11008). Fluorescent dyes used: Hoechst 33342 *1:500* (ThermoFisher, 62249) and Octadecyl Rhodamine B (R18) *1:1000* (ThermoFisher, O246).

### G. Image Analysis

Metrics describing spheroid area and circularity were quantified using Image J (version 1.51) software. In Image J, circularity is defined as (4π x area) / perimeter^2^ which ranges from zero to one, where one represents a perfect circle. Branch length was manually quantified as the length from the boundary between the branch and spheroid to the tip of the branch. Feature sizes less than 40 μm were excluded from analysis. The total number of branches, including all primary, secondary, and tertiary branches, was manually counted.

Immunofluorescence intensity data was analyzed via a custom pipeline developed in *CellProfiler* v4.2.6, an open-source segmentation algorithm designed to automate cellular phenotypic analysis^72^. To determine regional variations in immunostaining intensity, objects were first identified via the *IdentifyPrimaryObjects* module, and then subsequently a *MeasureObjectIntensityDistribution* module was employed to enable automatic and equidistant segmentation and analysis of the tip, body, and branch regions of the spheroids. This pipeline was conducted separately for all β1 integrin, phosphorylated FAK, and nuclei images. All phosphorylated FAK and β1 integrin intensity values were normalized against the intensity of the nuclei stain in each region.

To quantify the orientation of collagen fibers, an open-source quantitative tool was employed^73^. Briefly, images were imported into CurveAlign (4.0) and “CT” fiber analysis method via “Tiff Boundary” was selected. Branches were manually segmented, and ROIs were incorporated to delineate collagen fiber orientations along the branch axis and adjacent to the branch axis. This method of analysis has been previously reported^40^. The program generated a CSV file with all the orientation angles it tracked. To eliminate bias from the total number of collagen fibers counted, the data was binned and then divided by the total number of fibers found per each ROI. Polar plots were generated via a custom MATLAB script.

### H. Matrix Displacement Analysis

To generate the matrix displacement maps, bead channel images were generated into a stack and corrected for drift using the ImageJ plugin Linear Stack Alignment with SIFT. Then, the ImageJ particle image velocimetry (PIV) plugin was implemented using a cross-correlation window of 64 pixels and 32 pixels to yield 16-pixel displacement vectors, per previous established protocols^70,71^. This PIV analysis generated a vector field of matrix displacements, which was then imported into the ImageJ PIV plot plugin to generate magnitude and vector displacement plots, with a manually set vector scale threshold. Brightfield images of the spheroid were overlayed onto the matrix displacement field using ImageJ.

For quantification of bead displacement, Imaris 10.2.0 was used. The Track Spots algorithm was selected to track the identified spots as objects over several sequential frames in the time series dataset. ROIs were used to delineate regional differences in quantifications. The reference frame was manually oriented so that the y-axis was aligned toward the spheroid. The Autoregressive Motion algorithm was implemented to predict the location and direction of the spot based on the previous frame. The hyperparameters were selected to include a maximum distance of 5 μm to reduce errors in tracking and to ensure the predicted future position of a spot was reliably based on its previous position. Additionally, the max gap parameter was set to 3 μm to reduce the risk of incorrect spot connections across frames. The position data was exported as a .csv file, and a custom MATLAB script was written to group track IDs and quantify the aggregated displacement over time. Additionally, the script calculated the displacement delta over time for each track. The average of all beads in at least 15 spheroids across 3 replicates were analyzed to quantify the mean displacement along the branching axis.

### I. Inhibition Studies

Pharmacological inhibitors were added to the growth medium one hour after hydrogel gelation. Media with inhibitors was changed every two days. The inhibitors used were: 50 μM NSC 23766 (Tocris Bioscience, Rac1 inhibitor), 50 μM Blebbistatin (Cayman Chemical, non-muscle myosin II inhibitor), and 10 μM PF 573228 (Cayman Chemical, FAK inhibitor). Brightfield images were collected on day seven of culture and morphologically analyzed via methods previously described.

### J. Modulation of hyperosmotic and hypoosmotic pressure

Hyperosmotic stress was induced by adding various concentrations of PEG 400 (TCI America) to the culture medium at final concentrations of 0% wt/vol (0 mOsm/L), 1.5% wt/vol (37.5 mOsm/L) and 3% wt/vol (75 mOsm/L), which correspond respectively to osmotic pressure increases of 0 kPa, 92 kPa, and 197 kPa. Hypoosmotic stress was induced by replacing 20% or 40% of the culture medium with deionized water. Osmotic pressures were calculated via an empirically derived formula, where c represents the wt/vol concentration of PEG 400 in media (Equation 3)^74,75^. In osmotic pressure experiments where the culture medium remained unchanged throughout the experiment, hypertonic and hypotonic media was added on day zero, one hour after matrix gelation. Brightfield images of spheroids were collected on day seven and analyzed using methods previously described. In dynamic osmotic pressure experiments, hypertonic media was either: (1) added on day zero and replaced with standard growth medium on day four, or (2) added on day four following four days of culture in standard growth medium. Brightfield images were taken on days zero, four, and seven, and images were analyzed via methods previously described.

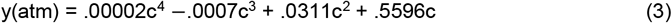

### K. Statistical Analysis and Reproducibility

All measurements were performed on at least three biological replicates from separate experiments. The sample size and statistical test performed for each experiment are indicated in the figure legends. Statistical analyses were all performed using GraphPad Prism 9.1.0. Specific tests for statistical significance are described in the figure captions.

## Acknowledgments

Funding for this project was provided by a National Science Foundation Graduate Research Fellowship (No: 2139319) to D.I.W., a National Science Foundation Data Driven Biology Grant (No. DGE-2125644) to D.I.W., a Crowe Family Fellowship to J.W.M., and a California Institute of Regenerative Medicine fellowship to A.S. (EDUC4-12821). The authors acknowledge the use of the Neuroscience Research Institute and the Department of Molecular, Cellular, and Developmental Biology Microscopy Facility supported by the University of California, Santa Barbara.

