## Supplementary Data for "Extracellular matrix viscoelasticity regulates mammary branching morphogenesis"

**Supplementary Video 1: Representative MCF10A spheroid in a fast stress relaxing matrix ($t_{1/2} \approx$ 100 s) undergoing isotropic expansion.** Time-lapse imaging shows brightfield channel overlaid with the fluorescent bead channel (green), taken 48 hours after encapsulation. Frames were acquired every 10 minutes for over 16 hours.

**Supplementary Video 2: Representative MCF10A spheroid in a slow stress relaxing matrix ($t_{1/2} \approx$ 1200 s) undergoing branching morphogenesis.** Time-lapse imaging shows brightfield merged with a fluorescent bead channel (green), taken after 48 hours after encapsulation. Frames are taken every 10 minutes for over 16 hours.

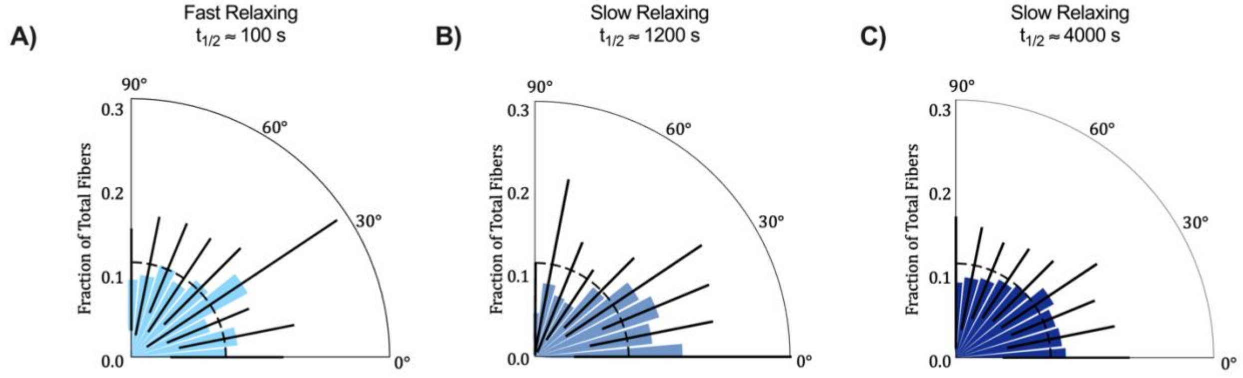


**Supplementary Figure 1: Collagen fiber alignment transverse to branching axis is randomly aligned regardless of matrix stress relaxation.** (a) Quantification of relative orientation angle transverse to the branching axis in the $t_{1/2} \approx$ 100 s matrix, (b) $t_{1/2} \approx$ 1200 s matrix, and (c) $t_{1/2} \approx$ 4000 s matrix. Collagen alignment was quantified via *CurveAlign*. n = 9 images from 3 hydrogels per condition. Black bars represent SD for each bin.

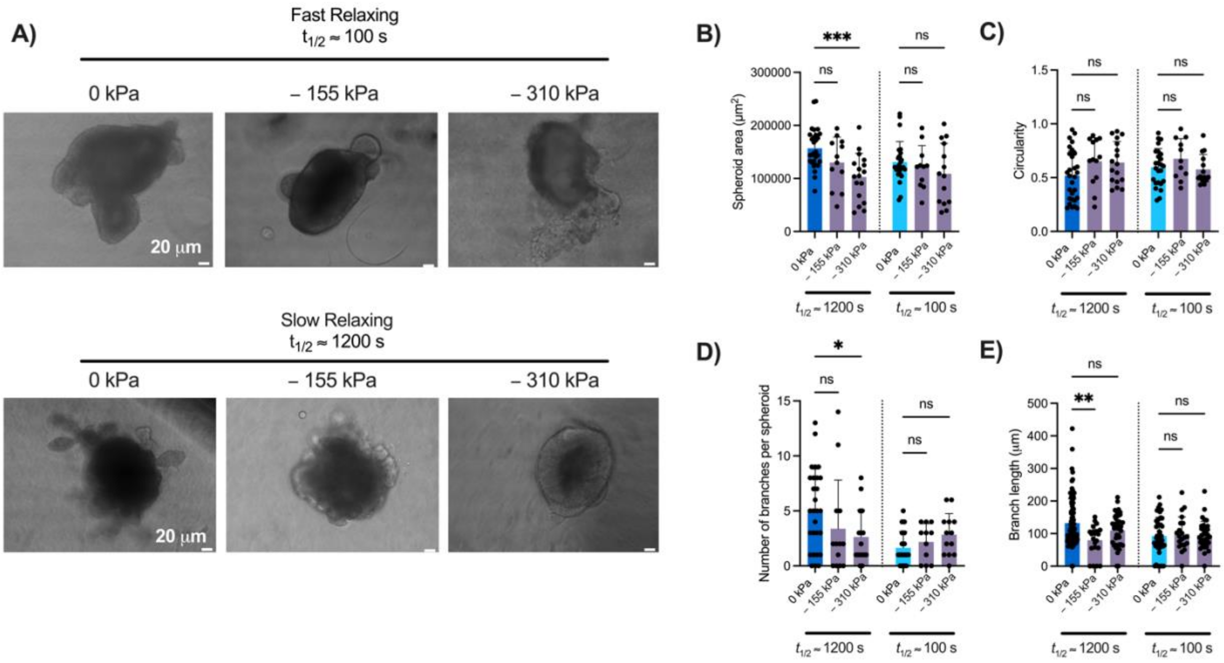


**Supplementary Figure 2: Hypoosmotic stress does not promote mammary branching.** (a) Brightfield images of MCF10A spheroids in various stress-relaxing conditions ($t_{1/2} \approx$ 100 s, $t_{1/2} \approx$ 1200 s) under different hypoosmotic stresses following 7 days of culture. (b) Quantification of MCF10A cross-sectional spheroid area under varied hypoosmotic pressure for 7 days. (c) Circularity is not significantly different in slow or fast relaxing matrices when spheroids are subjected to hypotonic media. (d) Quantification of branch length demonstrates that spheroids branch to a lesser extent in slow relaxing matrices when exposed to – 155 kPa. (e) Total branches produced per spheroid decreases when exposed to – 155 kPa in slow relaxing matrices, and there is no significant difference in fast relaxing matrices. Statistical significance calculated using one-way ANOVA with Šídák's multiple comparison tests: p*** < .001, p**<.01,*p<.05 and n.s. = not significant. n = 5-15 images from 3 independent experiments. All data are represented as mean +/- SD.

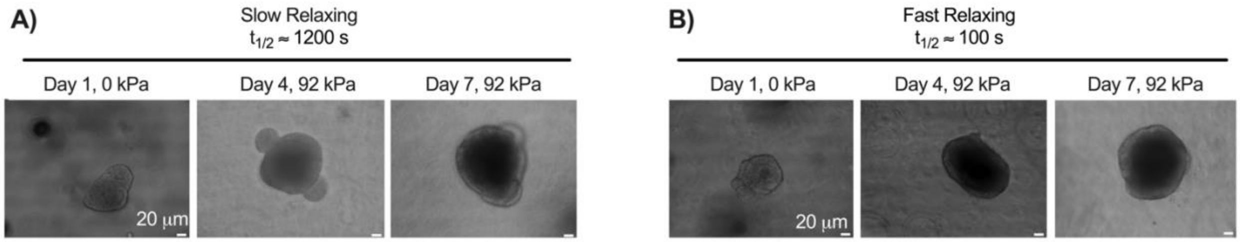


**Supplementary Figure 3: Hyperosmotic stress dynamically impedes mammary branching.** (a) Representative brightfield images of MCF10A spheroids in slow stress relaxing conditions ($t_{1/2} \approx$ 1200 s) and (b) fast stress relaxing conditions ($t_{1/2} \approx$ 100 s) after osmotic pressure (92 kPa) was applied on Day 4 in culture.

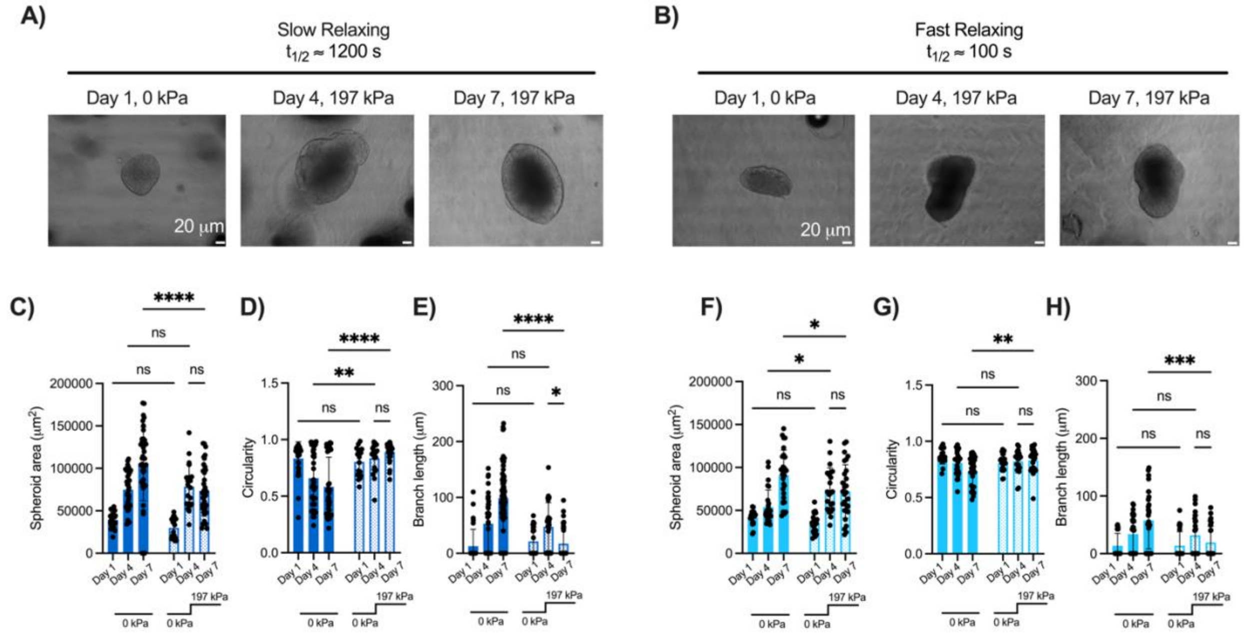


**Supplementary Figure 4: Dynamic changes to higher hyperosmotic stresses impair mammary epithelial branch formation and growth in slow stress relaxing matrices.** (a) Representative brightfield images of MCF10A spheroids in slow relaxing conditions ($t_{1/2} \approx$ 1200 s) and (b) fast stress relaxing conditions ($t_{1/2} \approx$ 100 s) after higher osmotic pressure (($\triangle P =$ 197 kPa) was applied on Day 4 in culture. (c) MCF10A spheriods are unable to resume growth in slow stress relaxing matrices and (f) fast stress relaxing matrices after osmotic pressure has been applied for 4 days. (d,e) Circularity is enhanced and branch length decreases when subjected to higher osmotic stresses in both slow and (g,h) fast relaxing conditions. Statistical significance calculated using one-way ANOVA with Šídák's multiple comparison tests: ****p < .0001, p*** < .001, p**<.01,*p<.05 and n.s. = not significant. n = 5-15 images from 3 independent experiments. All data are represented as mean +/- SD.

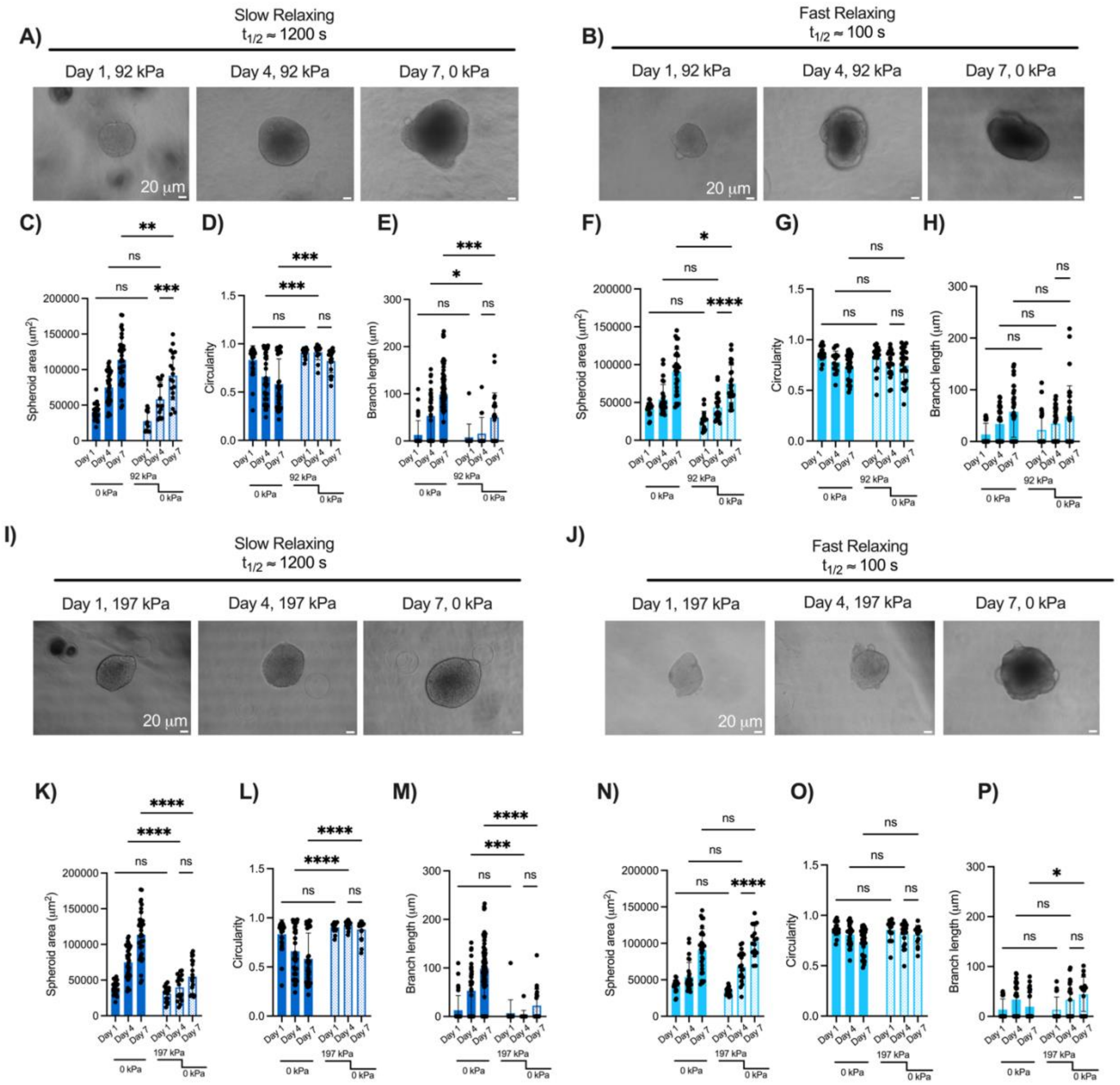


**Supplementary Figure 5: MCF10A cells are able to resume growth, but not branching, following removal of hyperosmotic stress**. (a) MCF10A spheroids are able to resume growth in slow stress relaxing matrices and (b) fast stress relaxing matrices after osmotic pressure ($\triangle P = 92$ kPa) has been applied for 4 days. (c-f) Quantification of MCF10A spheroid area demonstrates MCF10A spheroids can resume growth in both slow and fast relaxing matrices after osmotic pressure is alleviated. (g) Circularity is significantly enhanced in slow stress relaxing matrices following removal of hyperosmotic stress. (h) Branch MCF10A spheroids are unable to resume significant branching after being subjected to osmotic pressure in slow relaxing matrices. There is no significant difference in circularity or branch length once hyperosmotic pressures have been removed from fast relaxing matrices. (i) Representative brightfield images of MCF10A spheroids in slow relaxing conditions ($t_{1/2} \approx 1200$ s) and (j) fast stress relaxing conditions ($t_{1/2} \approx$ 100 s) after higher osmotic pressure ($\triangle P = 197$ kPa) was released on Day 4 in culture. (k) In slow relaxing matrices, cross-sectional area, (l) circularity, and (m) branch length significantly decrease compared to the control after osmotic pressure has been applied and released. (n) In fast relaxing matrices, there is no significant difference in cross-sectional area nor (o) circularity after osmotic pressure has been applied and released. (p) MCF10A spheroids in fast relaxing matrices can resume branching following the release of osmotic stress. Statistical significance calculated using one-way ANOVA with Šídák's multiple comparison tests: ****$p < .0001$, p*** < .001, p**<.01,*$p<.05$ and n.s. = not significant. n = 5-15 images from 3 independent experiments. All data are represented as mean +/- SD.
